# Preclinical and Toxicology Studies of BRD5529, a Selective Inhibitor of CARD9

**DOI:** 10.1101/2021.11.19.469250

**Authors:** Theodore J. Kottom, Kyle Schaefbauer, Eva M. Carmona, Eunhee S. Yi, Andrew H. Limper

## Abstract

**Background:** Exuberant inflammation during *Pneumocystis* pneumonia leads to lung injury. CARD9 is a central mediator of inflammatory signaling mediated by C-type lectin receptors. CARD9 inhibitor BRD5529 has been shown to be an effective *in vitro* inhibitor of *Pneumocystis* β-glucan-induced proinflammatory signaling and downstream TNF-alpha production, suggesting its viability as a candidate for preliminary drug testing as an anti-inflammatory agent in the rodent Pneumocystis pneumonia model (PCP).

**Methods:** To assess for potential toxicity, mice were injected intraperitoneally (IP) daily either with vehicle or BRD5529 at 0.1 mg/kg or 1.0 mg/kg for two weeks. Mouse weights were taken daily. At day 14, mice were euthanized, weighed, and analyzed by flexiVent™ for lung stiffness. Lungs, liver, and kidney were then harvested for H&E staining and pathology scoring. Lung samples were further analyzed for proinflammatory cytokines via ELISA and extracellular matrix generation via quantitative PCR (q-PCR). Blood collection postmortem was performed for blood chemistry analysis.

**Results:** BRD5529 at both doses of IP administration resulted in no significant changes in daily or final weight gain. Analysis of lung stiffness by flexiVent™ showed no significant differences between the control or treated groups. Furthermore, ELISA results for proinflammatory IL-1 Beta, IL-6, and TNF-alpha showed no major differences in the respective groups. qPCR analysis of extracellular matrix transcripts collagen type I, alpha 1 (*Col1a1*) and fibronectin (*Fn*) were statically similar as well in the treated and control groups. Examination and pathology scoring of H&E slides from lung, liver, and kidney from the each of the mice in all groups and subsequent pathology scoring showed no significant change. Blood chemistry analysis revealed similar, non-significant patterns.

**Conclusions:** BRD5529 in our initial general safety and toxicology assessments displayed no inherent safety concerns in the analyzed parameters. These data support broader *in vivo* testing of the inhibitor as a timed adjunct therapy to the deleterious proinflammatory host immune response often associated with anti-*Pneumocystis* therapy.

## 1. Introduction

Caspase recruitment domain-containing protein 9 (CARD9) is a central mediator downstream of C-type lectin receptors (CLRs) that is vital for microbial pathogen proinflammatory host immune response and organism burden control [1]. CARD9 is highly expressed in myeloid cells and shown to be particularly important in fungal infections [2]. Others have demonstrated CARD9 pathway intervention with the chemical CARD9 inhibitor BRD5529 can directly mimic a protective variant of the protein and may provide therapeutic benefit with those with inflammatory bowel disease [3]. Recently, we have shown that preincubation of BRD5529 with RAW macrophages prior to the application of proinflammatory β-glucans from the lung pathogen *Pneumocystis* spp., results in substantial reduction in downstream CARD9 proinflammatory signaling and subsequent TNF-alpha release, suggesting that timed therapeutic intervention during or after anti-*Pneumocystis* treatment, may greatly improve the deleterious effects on the host caused by organism killing and release of proinflammatory carbohydrates [4]. The purpose of this study was to evaluate the short-term administration of CARD9 inhibitor BRD5529 in mice via IP administration and address potential detrimental responses to the inhibitor via physiological, inflammatory, and toxicological analysis. These data demonstrate the safety of BRD5529 and support broader clinical development of the CARD9 inhibitor for *in vitro* administration in the PCP mouse treatment model as a therapeutic tool to treat PCP host inflammation during anti-*Pneumocystis* treatment.

## 2. Methods

### 2.1. Animals

Equal numbers of male and female C57BL/6 mice (The Jackson Laboratory) at 10- 12 weeks of age were used for all experiments. All animal procedures were performed in accordance with the Laboratory Animal Welfare Act, the Guide for the Care and Use of Laboratory Animals Welfare Act, and the Mayo Clinic Institutional Animal Care and Use Committee (IACUC) (Approval number: A00005722-20).

### 2.2. Administration of BRD5529

BRD5529 was obtained for Sigma Aldrich. Due to the lack of solubility of the inhibitor in water or saline, the inhibitor was prepared with 1% Methocel™ [5]. Intraperitoneal treatment (100 μl) with 1% Methocel™ (vehicle, control mice group) or the indicated concentration (mg/kg) of the BRD5529 inhibitor in Methocel™ was initiated on day 0 and subsequentially every day for 14 days. At day 14, mice were sacrificed, and subsequent analysis performed as described below.

### 2.3. Flexivent™ analysis

FlexiVent™ analysis was performed at described previously [6].

### 2.4. ELISA determination of cytokine release

Cytokines were analyzed from total lung homogenates. ELISA kits to measure mouse IL-1 Beta ,IL-6, and TNF-alpha were purchased from Thermo Fisher Scientific.

### 2.5. Quantitative polymerase chain reaction analysis

To extract RNA from mouse lung, tissue was lysed and homogenized with Buffer RLT Plus (supplied with the RNeasy^®^ Plus Mini Kit; Qiagen). The lysate was passed through a genomic DNA eliminator spin column, ethanol was added, and the sample was applied to a RNeasy MinElute spin column according to the manufacturer’s instructions. An iScript™ Select cDNA synthesis kit (Bio-Rad) was used for reverse transcription using oligo (dT) primers and random hexamer primer mix. A SYBR green PCR kit (Bio-Rad) was used for quantitative real-time PCR and was performed and analyzed on a CF96 Touch™ Real-Time PCR Detection System (Bio-Rad). The sequences of the primer pairs are listed in Supplementary Table 1.

### 2.6. Biochemical analysis

For blood chemistry analysis, serum was analyzed with the Piccolo Xpress™ Chemistry Analyzer according to the manufacturer’s instructions.

### 2.7. Histology analysis

For histological analysis, lung, liver, and kidney samples were fixed in 10% neutral formalin. Paraffin embedding and staining were performed at the Mayo Clinic Histology Core, Scottsdale, AZ. Sections (5 μm) were stained with Hematoxylin and eosin (H&E) and graded blindly for the extent of organ inflammation by a Mayo Pathologist. The sections were scored as follows: 1+, mild perivascular aggregates; 2+, heavy perivascular aggregates; 3+, mild alveolar aggregates; 4+, alveolar exudate and heavy alveolar aggregates; and 0, normal. These scores were based upon grading of the entire organ surface area present on the slide section.

### 2.8. Statistical Analysis

For multigroup data, initial analysis was first performed with analysis of variance (ANOVA) to determine overall different differences. If ANOVA indicated overall differences, subsequent group analysis was then performed by 2-sample unpaired Student’s *t*-test for normally distributed variables. Evaluation of data was conducted using Prism 9 for macOS, version 9.1.0 (GraphPad). Values of *p* < 0.05 were considered significant.

## 3. Results

### 3.1. BRD5529 IP administration resulted in no significant weight loss

### 3.2. Static lung compliance after BRD5529 IP administration

To measure lung function in mice after 14 days of IP administration of vehicle, or BRD5529 at 0.1 mg/kg or 1.0 mg/kg we used the flexiVent™ apparatus. Daily IP injections of all three conditions for 14 days resulted in no significant increases in lung static compliance between the three groups tested (**Fig 2**).

**Fig. 1.**
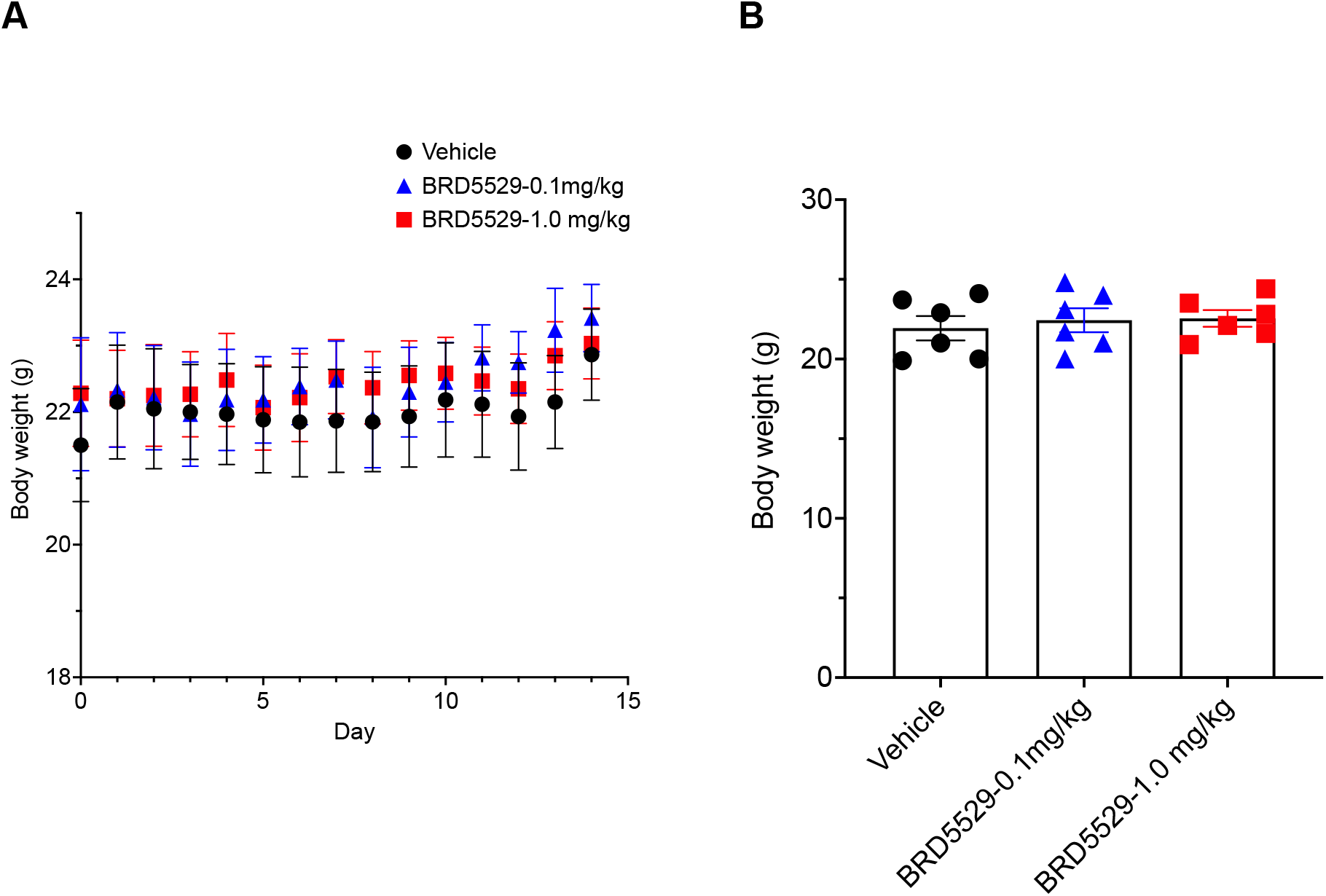
Effects of mouse body weight with IP administration daily of BRD5529. (A) Shows the daily weight changes in the vehicle control versus the 0.1 mg/kg and 1.0 mg/kg doses of IP administered BRD5529 daily for 14 days. (B) Shows the final weights for the 3 groups after 14 days. (n = 6 mice/group). No significant weight changes were noted between the three groups.

**Fig. 2.**
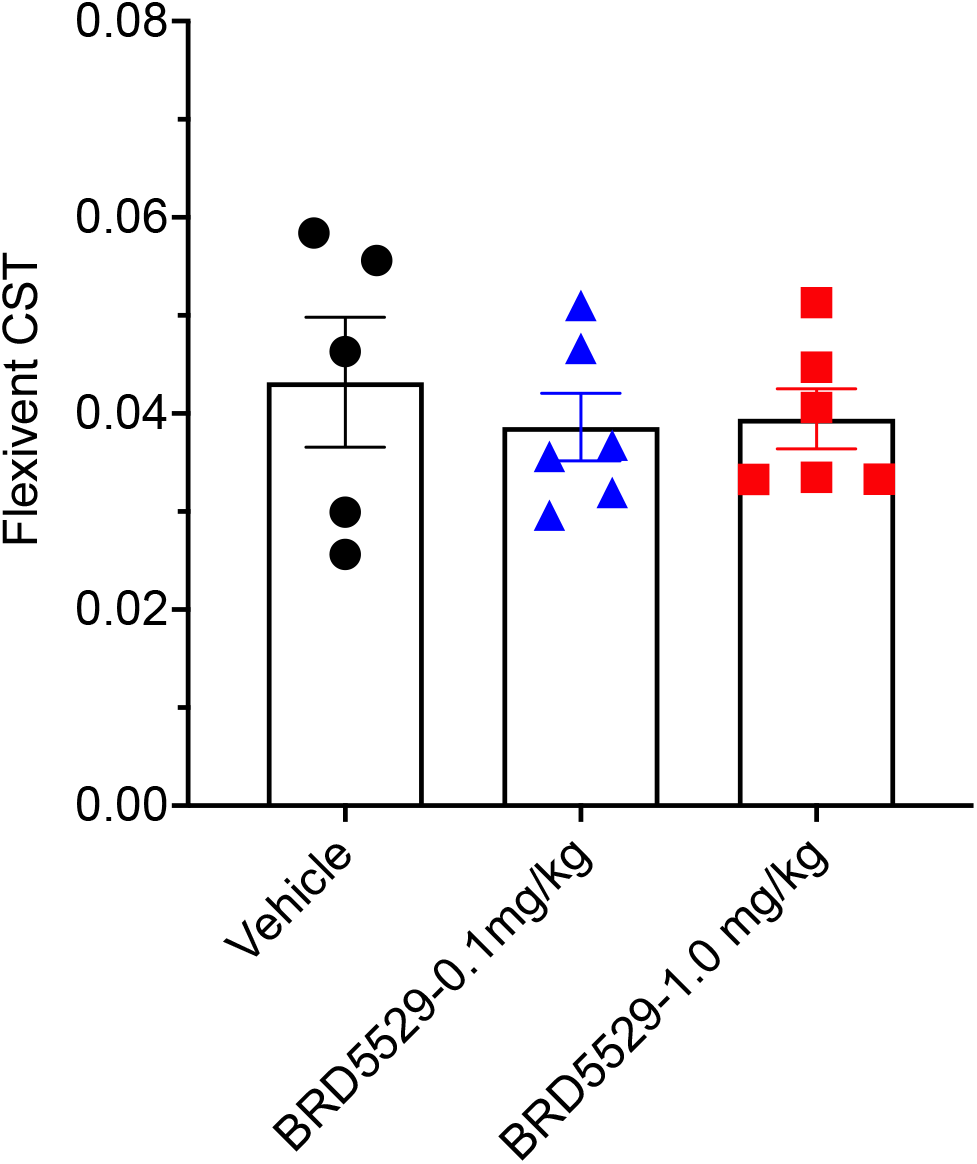
CARD9 inhibitor BRD5529 effects on lung compliance. Lung quasi-static compliance (CST; reflects the intrinsic elastic properties of the lung and chest at rest). (n = 6 mice/group). No significant changes in CST were noted between the three groups.

### 3.3. Measurements of lung cytokines in BRD5529 IP administrated and control groups

The production of inflammatory cytokines IL-1 Beta, IL-6, and TNF-alpha were measured in whole lung lysates and as shown in (**Fig. 3A-C**). No significant alterations were noted from the vehicle control.

**Fig. 3.**
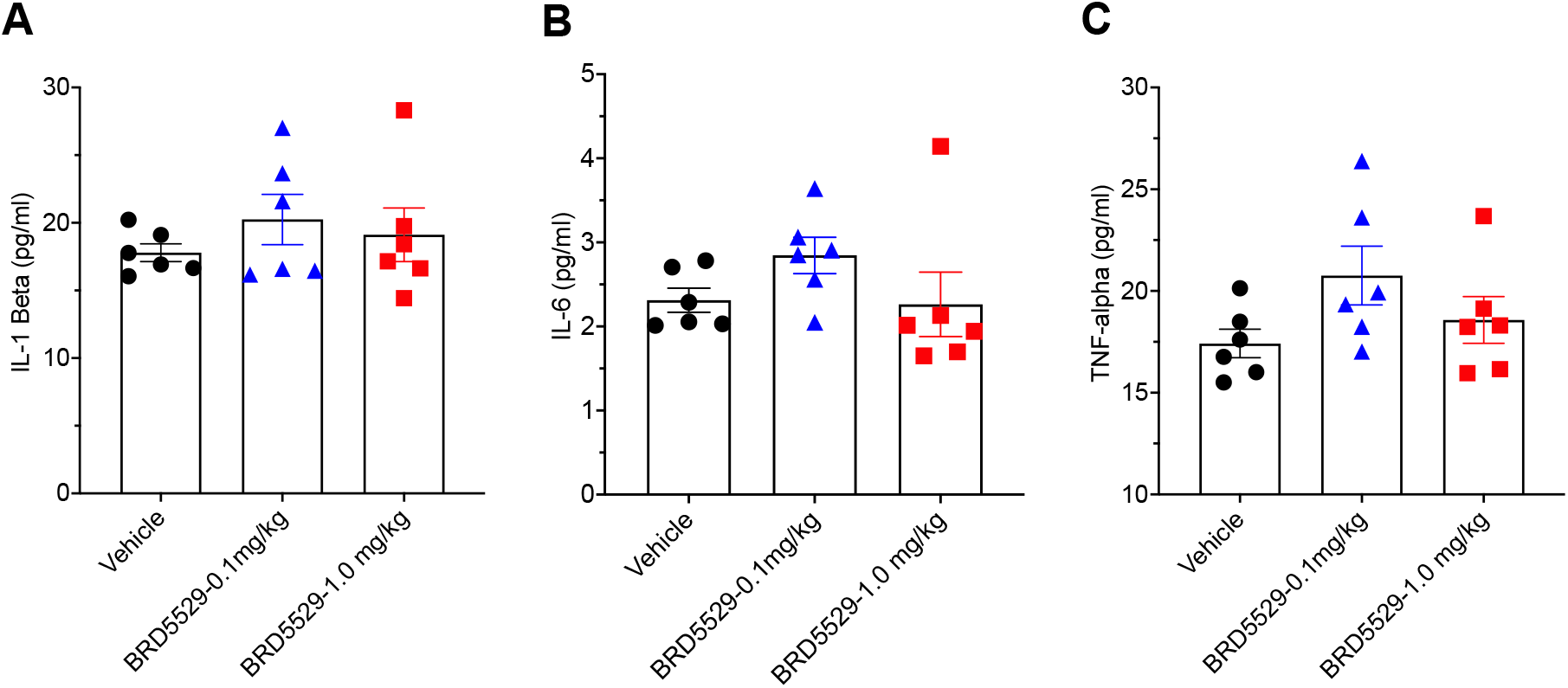
CARD9 inhibitor BRD5529 effects on lung proinflammatory cytokine production. (A) IL-1 Beta, (B) IL-6, and (C) TNF-alpha production was measured from total lung lysates from day 14 of the experiment. (n = 6 mice/group). No significant differences were noted between the groups.

### 3.4. Analysis of mRNA extracellular matrix generation

qPCR was implemented to determine the levels of mRNA expression of Collagen Type Alpha 1 Chain (*Col1a1*) and Fibronectin (*Fn*), both extracellular matrix related genes used as markers for profibrotic development [7]. Beta 2 Microglobulin (*B2M*) was used as a housekeeping gene. As shown in (**Fig. 4**A-B), no significant differences were noted in the 3 groups at day 14 in the respective lung samples.

**Fig. 4.**
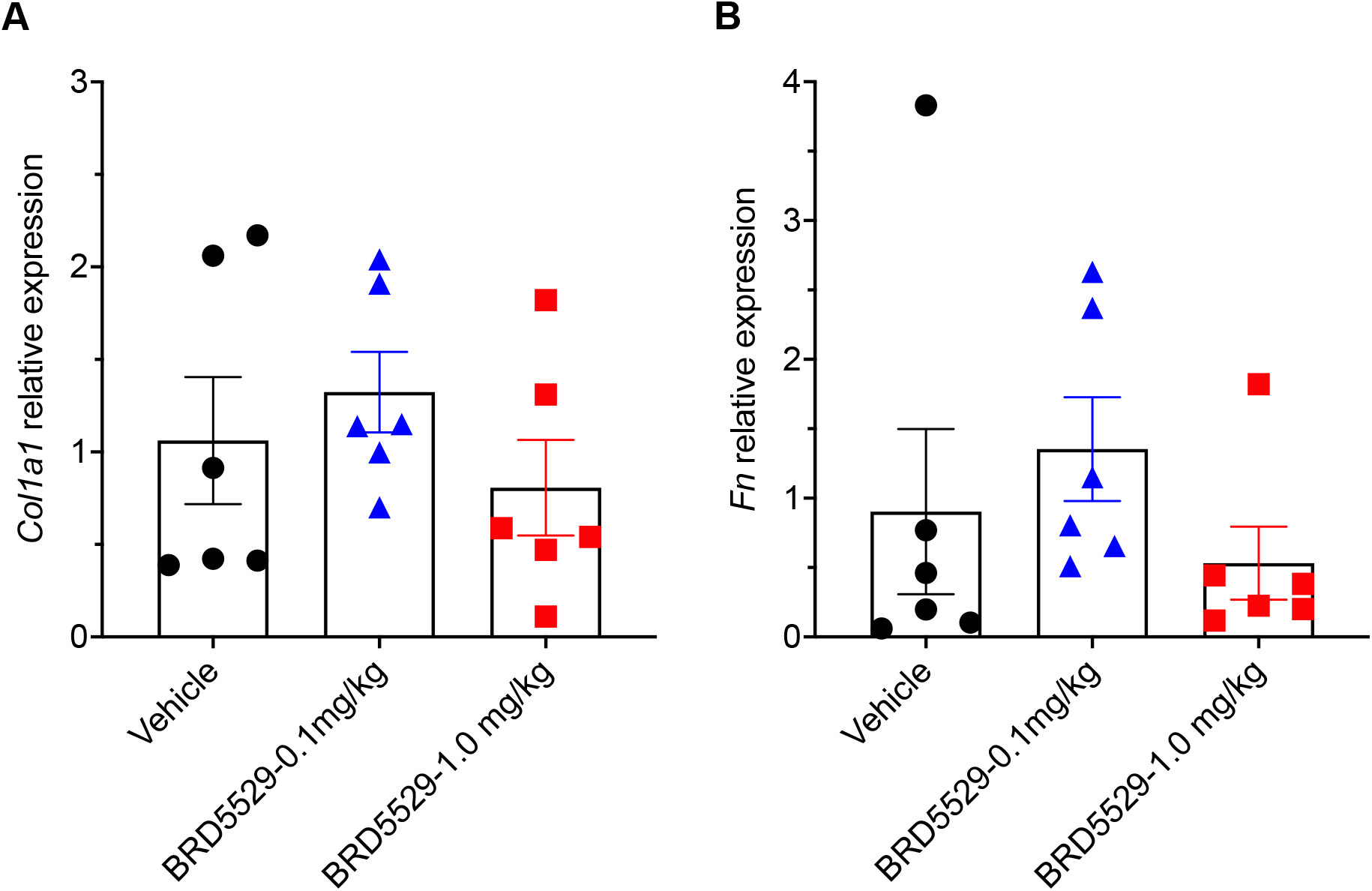
Quantitation of *Col1a1* and *Fn* mRNA in total lung RNA after vehicle and BRD5529 administration for 14 days. Ratios of (A) *Col1a1* and (B) *Fn* to *B2M* in total lung RNA. (n = 6 mice/group). No significant differences were noted between the groups.

### 3.5. Serum chemistry data

Complete group mean serum chemistry data from data 14 are presented in (**Fig. 5**A-N). There were no noteworthy changes in any either BRD5529 dose groups from the vehicle control.

**Fig. 5.**
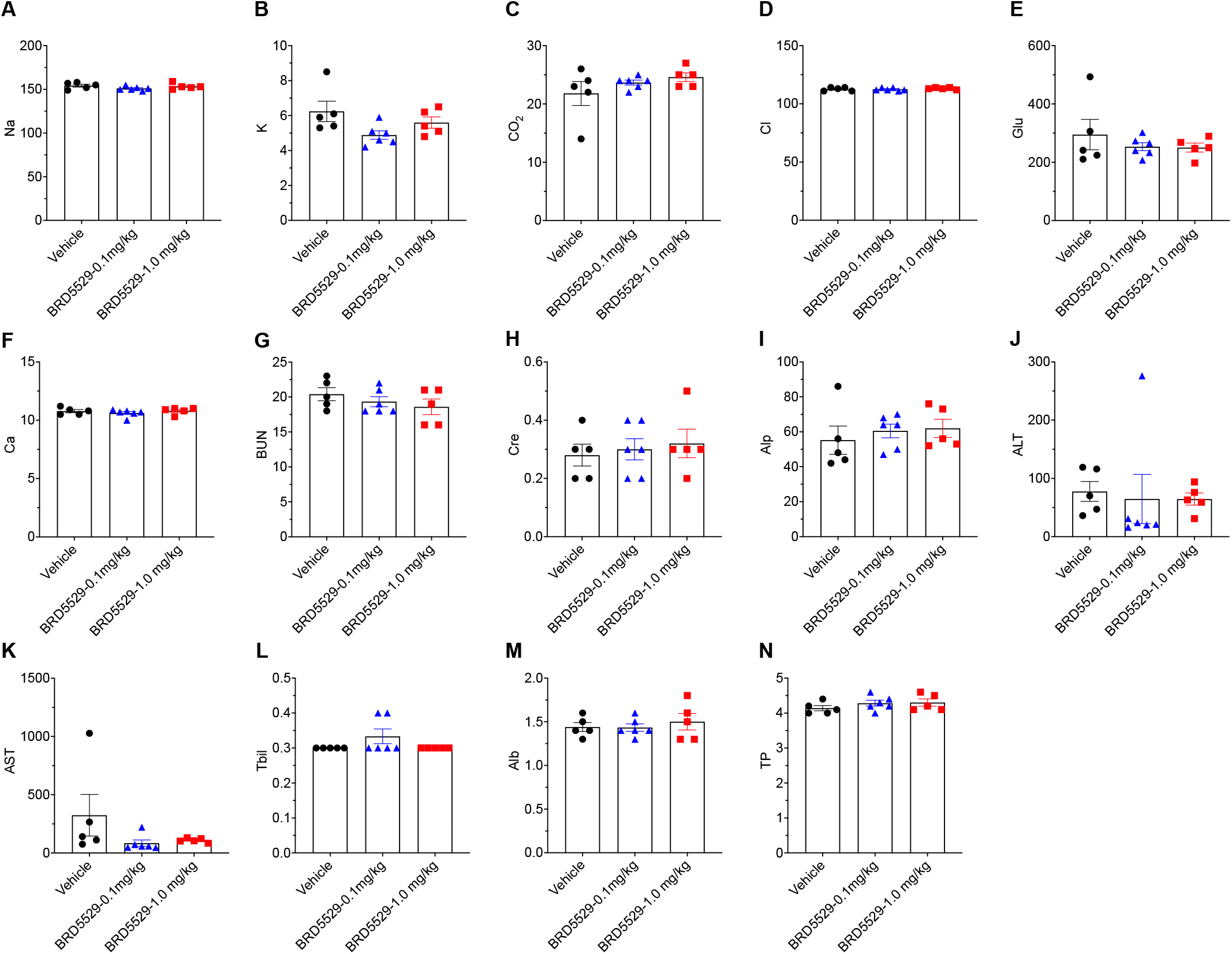
Serum chemistry parameters. Na, sodium (mmol/L); K, potassium (mmol/L); CO_2_, carbon dioxide (mmol/L); Cl, chloride (mmol/L); Glu, glucose (mg/dL); Ca, Calcium (mg/dL); BUN, blood urea nitrogen (mg/dL); Cre, creatine (,g/dL); Alp, alkaline phosphatase (U/L); ALT, alanine aminotransferase (U/L); Tbil, total bilirubin (mg/dL); Alb, albumin (g/dL); TP, total protein (g/dL). (n = 5-6 mice/group). No significant differences were noted between the groups.

**Fig. 6.**
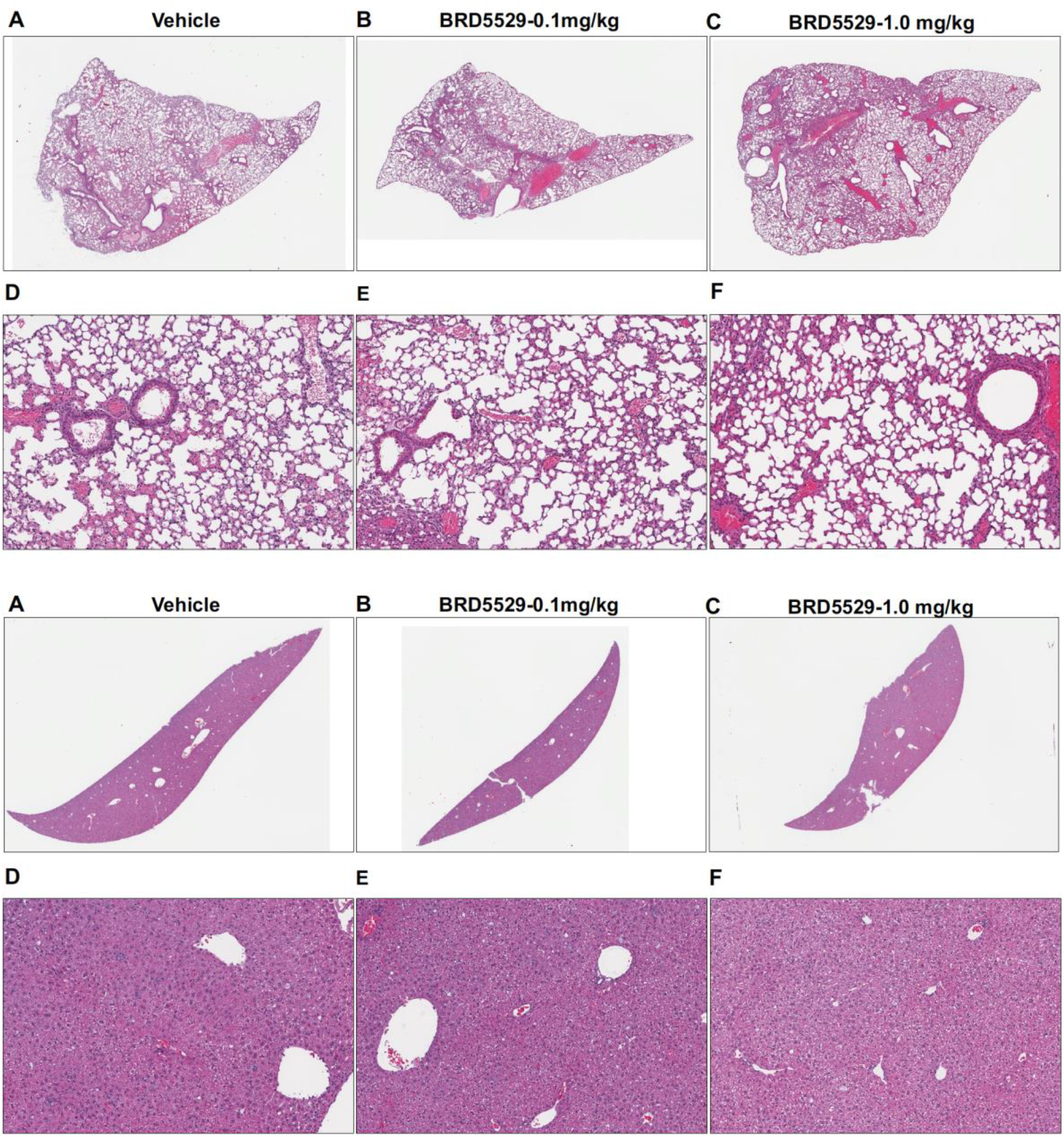
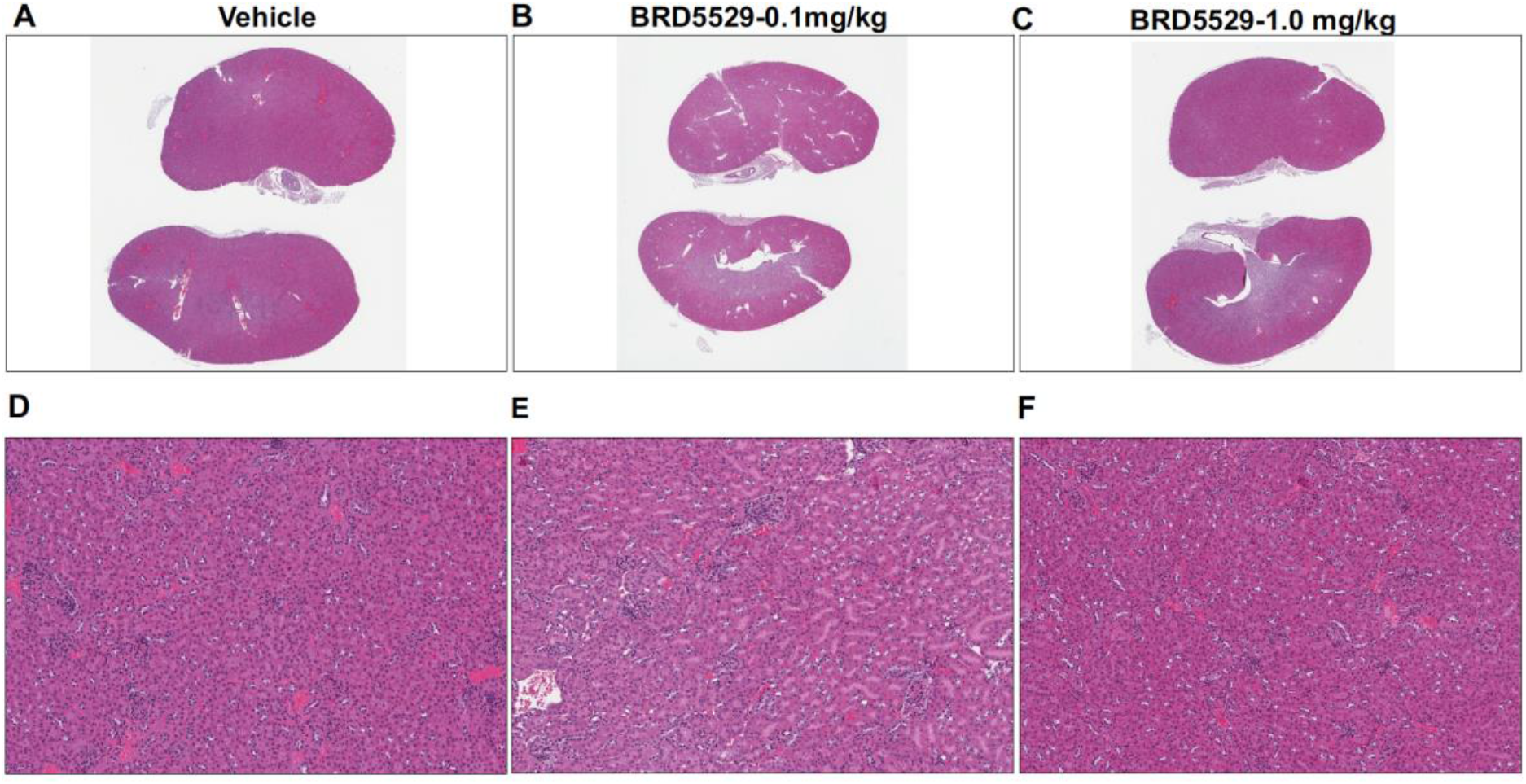
Lung, liver, and kidney histopathology of 14-day IP treated vehicle or BRD5529 CARD9 inhibitor. Hematoxylin and eosin (H&E) staining was performed on sections of lung (top two panels), liver (middle two panels), and kidney (bottom two panels) from mice in all groups. (A) Vehicle control. (B) BRD5529 at 0.1mg/kg. (C) BRD5529 at 1 mg/kg. No histological changes were present in lung, liver, and kidneys in the vehicle and BRD5529 doses tested.

### 3.6. Histology analysis

Histologic examination of all samples from lung, liver, and kidney from both BRD5529 treated- and vehicle groups did not reveal any abnormality and all organs appear normal (score 0 for all parameters).

## 4. Discussion

Fungi are major contributors to opportunistic infections in those with advanced HIV. Pneumocystis jirovecii pneumonia (PCP), caused by *Pneumocystis jirovecii* is among the most common pathogens in AIDS populations across the globe. Implementation of highly active antiretroviral therapy (HAART) has decreased the overall incidence of PCP, but the ability to receive this treatment is limited and cases of PJP requiring hospitalization is still quite high [8].

The most current anti-*Pneumocystis* therapy, trimethoprim-sulfamethoxazole (TMP-SMX), has proven to an effective antimicrobial combination to treat *Pneumocystis jirovecii* pneumonia (PJP). Although effective, the exuberant inflammatory response following fungal cell death, via the exposure of newly exposed β-glucans can prove highly detrimental to the host [9-11]. Indeed, when corticosteroids are utilized in HIV patients with moderate-severe PJP, there is a significant decrease in mortality and morbidity noted [12]. In the non-HIV patient, data on adjuvant corticosteroids is less clear and no consensus has been determined. Even though steroids may be beneficial to the patient in these settings, there are still both short-term (co-infections, hyperglycemia) and long term (myopathy and osteoporosis) that need to be considered [13].

Therefore, other adjunct therapies should be considered in PJP. Recently, we have demonstrated that *in vivo*, macrophages pre-incubated with the caspase recruitment domain-containing protein 9 (CARD9) inhibitor BRD5529 have significant reductions in their ability to generate proinflammatory signaling and downstream TNF-alpha production upon stimulation with Pneumocystis β-glucans [4]. These results lead us to hypothesize that BRD5529 may be used *in vivo* as an adjunct therapy similar to corticosteroids [4]. As part of the development toward human clinical application, a thorough preclinical assessment must be conducted to evaluate safety and potential toxicity of the BRD5529 CARD9 inhibitor, and to the best of our knowledge this is the first study conducted to examine this. The preclinical profile presented here suggests thar BRD5529 could meet these early criteria.

In this study we gave mice BRD5529 either at 0.1 mg/kg or 1.0 mg/kg once a day for 14 days via IP administration. In acute toxicity studies, administration of BRD5529 appeared to be well tolerated in mice at both doses. Parameters such as weight loss, lung function, lung-specific proinflammatory response, lung extracellular matrix mRNA generation, blood toxicology analysis, and H&E histological examine of lung, liver, and kidney samples yielded no significant changes as compared to the vehicle control.

## 5. Conclusions

Based on these preliminary findings, the use of BRD5529 in the future employing *in vivo* mouse studies to determine whether the CARD9 inhibitor can be used to reduce the deleterious effects of the host proinflammatory in the PCP model seems safe and feasible.

## Abbreviations

CARD9: (Caspase recruitment domain-containing protein 9)
PCP: (*Pneumocystis* pneumonia)
IP: (Intraperitoneally)
(IACUC): Institutional Animal Care and Use Committee
CLR: (C-type lectin receptor)
(H&E): Hematoxylin and eosin
HAART: (highly active antiretroviral therapy)
(TMP-SMX): trimethoprim-sulfamethoxazole
(PJP): Pneumocystis jirovecii pneumonia

## CRediT authorship contribution statement

**Theodore J. Kottom**: Writing – original draft, drafted the manuscript, performed the experiments, performed data analysis. **Kyle Schaefbauer:** performed the experiments. **Eva M. Carmona**: Writing – review and editing. **Eunhee S. Yi**: Writing – review and editing, performed data analysis. **Andrew H. Limper**: Writing – review and editing, helped with data design and analysis.

## Supplementary Materials

**“Preclinical and Toxicology Studies of BRD5529, a Selective Inhibitor of CARD9” TJ Kottom, AH Limper, et al.**

**Supplementary Table 1.**
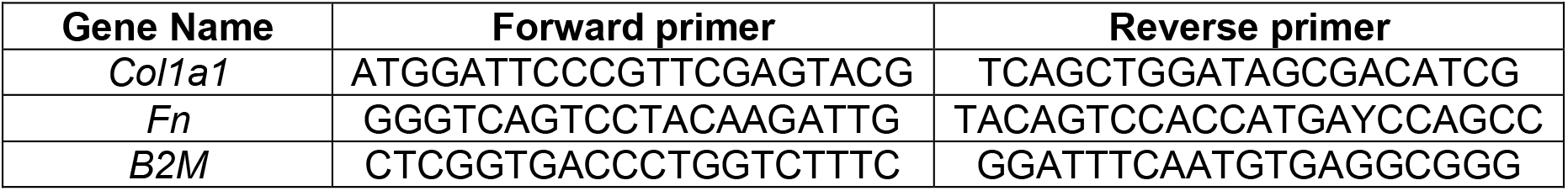

